# Alterations in odor hedonics in the 5XFAD Alzheimer’s disease mouse model and the influence of sex

**DOI:** 10.1101/2020.05.08.085043

**Authors:** Elizabeth R. Roberts, Amanda M. Dossat, María del Mar Cortijo, Patrik Brundin, Daniel W. Wesson

**Affiliations:** Department of Pharmacology & Therapeutics, University of Florida, 1200 Newell Dr., Gainesville, FL, 32610; Van Andel Institute, 333 Bostwick Ave NE, Grand Rapids, MI 49503

**Keywords:** Tg6799, olfaction, sex difference, valence, behavior

## Abstract

Olfactory impairments, including deficits in odor detection, discrimination, recognition, and changes in odor hedonics are reported in the early stages of Alzheimer’s disease (AD). Rodent models of AD display deficits in odor learning, detection, and discrimination – recapitulating the clinical condition. However, the impact of familial AD genetic mutations on odor hedonics is unknown. We tested 2-, 4-, and 6-months old 5XFAD (Tg6799) mice in the five-port odor multiple-choice task designed to assay a variety of odor-guided behaviors, including odor preferences/hedonics. We found that 5XFAD mice investigated odors longer than controls, an effect that was driven by 6-months old mice. Interestingly, this effect was carried by females in the 5XFAD group, who investigated odors longer than age-matched males. Upon examining behavior directed towards individual odors to test for aberrant odor preferences, we uncovered that 5XFAD females at several ages displayed heightened preferences towards some of the odors, indicating aberrant hedonics. We observed no impairments in the ability to engage in the task in 5XFAD mice. Taken together, 5XFAD mice, particularly 5XFAD females, displayed prolonged odor investigation behavior and enhanced preferences to certain odors. The data provide insight into hedonic alterations which may occur in AD mouse models, and how these are influenced by biological sex.

## Introduction

Alzheimer’s disease (AD) is a neurodegenerative disorder characterized by cognitive, motor, and sensory dysfunction. Olfactory dysfunction is a common feature of the AD prodrome, with patients displaying deficits in olfactory detection, discrimination, and recognition (Devanand et al., 2008; Doty, 2017; Murphy, 2019; Rey et al., 2018). The emotional responses to odors, known as hedonics, are also disturbed in persons with AD (Joussain et al., 2015). While alterations in odor hedonics have received less attention in AD research than manifestations in perceptual impairments, this is a significant issue since odor preferences influence food selection and ingestive behaviors (Murphy, 2008) which, if aberrant, might impact disease progression, and at a minimum, quality of life (Gopinath et al., 2012; Murphy, 2019).

Mouse models of AD may provide insight into the origins of aberrant odor preferences in the disease. These mouse models have already yielded clues into some mechanisms of olfactory dysfunction and how these deficits might relate to disease progression. For instance, the Tg2576 mouse model of AD, which over-expresses two familial mutations in the amyloid precursor protein gene (*APP*), displays impairments in short-term odor learning and odor discrimination (Wesson et al., 2010). These deficits are associated with alterations in neural activity in the olfactory bulb and piriform cortex – two important regions of the olfactory system (Wesson et al., 2011). Transient over-expression of mutant *APP* in solely the olfactory sensory neurons in the nasal epithelium is sufficient to impair olfactory perception (Cheng et al., 2011). While not well understood, it is possible that products of the *APP* gene, and/or other genes implicated in AD, impact olfactory system physiology and thus perturb perception (Morales-Corraliza et al., 2012; Wesson et al., 2010, 2011; Xu et al., 2015; Yoo et al., 2017). Whether animal models which express these pathologies have alterations in their odor hedonics is unknown.

We sought to investigate odor preferences in an AD mouse model. To address this, we used the 5XFAD (Tg6799) model which overexpresses five familial AD mutations and thus is considered an accelerated AD model (Oakley et al., 2006). The 5XFAD mouse model displays progressive AD pathology throughout the brain, starting at just two months of age (Oakley et al., 2006), which differentially burdens males and females (Bundy et al., 2019). This includes amyloid-β burden in some key olfactory structures, such as the olfactory bulb and piriform cortex (Oakley et al., 2006; Wang et al., 2017; Xiao et al., 2015). Further, this mouse model shows impairments in synaptic plasticity and cognition (Devi & Ohno, 2016; Kimura & Ohno, 2009; Oakley et al., 2006). Importantly, 5XFAD mice appear to exhibit normal olfactory detection abilities, as assayed in a sensitive psychophysical assay (Roddick et al., 2016). However, the possibility that despite intact sensitivity for odors, these mice may have aberrant tendencies to investigate odors and possibly altered hedonics to select odors, has not yet been investigated. In both humans and mice, the duration of time spent investigating odors reflects the attractiveness or preference of an odor reflective of its hedonics (Baum & Keverne, 2002; Mandairon et al., 2009). Therefore, we monitored the duration of time 5XFAD mice spent investigating odors in a five-port odor multiple-choice task, which is a sensitive and semi-automated assay of odor preferences (Jagetia et al., 2018). We did so grouping across all odors to test for general tendencies to investigate odors, and separately, within individual odors to explore possible alterations in hedonics. Our results show that 5XFAD mice, particularly 5XFAD females, display prolonged odor investigation behavior and enhanced preferences to certain odors.

## Materials & Methods

### Subjects

We used ninety-six 5XFAD/B6SJL-Tg6799 (5XFAD) mice and age-matched nontransgenic (NTg) littermates (2-, 4-, and 6-months of age; 48 males and 48 females). Breeder stock were obtained from The Jackson Laboratory (stock #006554; Bar Harbor, ME) and their offspring were raised and maintained at the University of Florida. Animals were genotyped using PCR of tail clip DNA upon weaning, and genotypes were confirmed with a second round of genotyping after completion of behavioral testing. All experiments were performed in accordance with the guidelines of the National Institutes of Health and were approved by the University of Florida Institutional Animal Care and Use Committee. From weaning onward, all animals were group-housed (2-5 per cage), placed on a reversed 12:12h light cycle, and allowed *ad libitum* access to food and water. Cohorts ranging from four to 16 mice were pseudo-randomly assigned to maximize diversity across sex, age, and genotype. All behavioral testing occurred during the light phase (0900:1800 hr).

### 5-port odor preference apparatus

We used the five-port odor multiple-choice behavioral apparatus as we previously described in detail (Jagetia et al., 2018). Briefly, this apparatus allows animals to freely investigate different 3D-printed odor ports and provides automated, millisecond-precise monitoring of nose-poking behavior towards odors by means of infrared photobeams. In this apparatus, animals gain access to the odor by nose-poking into a port with nose-pokes detected by an 880 nm infrared beam positioned at the entrance of the odor port. The odor ports were designed to scavenge odor out of the port, thereby requiring the animal to poke to gain access to the odor (Jagetia et al., 2018). On the testing day, 20 µL of liquid odor was dispensed into a flange cap (MOCAP, Park Hills, MO) and inserted into the bottom of a port. The onset, duration, and location (port identity) of nose-pokes were recorded by a microcontroller (Arduino; www.arduino.cc), digitized, and stored on a computer for subsequent analysis.

### Five-port odor multiple-choice preference testing

On acclimation and testing days alike, the mice were brought into the testing room where they were allowed to acclimate in their home cages prior to being placed in the testing chamber. The mice were allowed to explore the five-port chamber for increasing durations over three consecutive days (10, 20, and 30 minutes, respectively). During acclimation, clean plastic flange caps were placed in the bottom of the odor ports. Within two days following acclimation, 30 minute testing sessions were conducted over five consecutive days. The same five odors were presented every day, with each odor being placed in a different port each day to minimize the influence of spatial location on odor investigation (Jagetia et al., 2018). The apparatus was cleaned with 70% ethanol before onset of testing and after each mouse was removed from the chamber.

### Odors

Odors were purchased at their highest available purity (>97%) and diluted to 1 Pa in light mineral oil (Sigma Aldrich; St. Louis, MO) prior to placing in the flange caps / odor ports and included L-carvone, decanol, geraniol, guaiacol, and thioglycolic acid (Sigma Aldrich; St. Louis, MO). Odors in their stock concentration and diluted concentration were each stored under nitrogen to maintain purity. The panel of odors selected was based upon our prior work (Jagetia et al., 2018), wherein we found that [at least in a different background strain of mice] these odors elicited the widest assortment of odor preferences, with decanol and thioglycolic acid being investigated the least and most, respectively.

### Data Analysis

Individual animals’ behavior was evaluated by the duration of nose-pokes (*viz*., odor investigation), nose-poke number, and inter-poke interval. Statistical analyses were performed in Prism 8.0 (GraphPad Software; San Diego, CA), included mixed-effects models with Geisser-Greenhouse corrections and two-way analysis of variance (ANOVA), and were followed by Sidak’s post hoc tests where indicated. Two-tailed unpaired t-tests were used to determine effects of genotype within each sex and age group. Any values that were > 2 standard deviations beyond the group mean were classified as outliers and removed from the data set. Across variables there was a range of 0-3 outliers per group, represented similarly across all genotypes, ages, and sexes. Over testing days, mice robustly habituate to odors in this task (Jagetia et al., 2018) (see also **Figures 1 and 2**). Therefore, in order to investigate possible changes in odor investigation and odor preferences between genotypes (as well as sex and age effects), we restricted our primary analyses to the behavior from day 1 of testing since this allows for examination of animals’ response to odors when they are highest.

**Figure 1.**
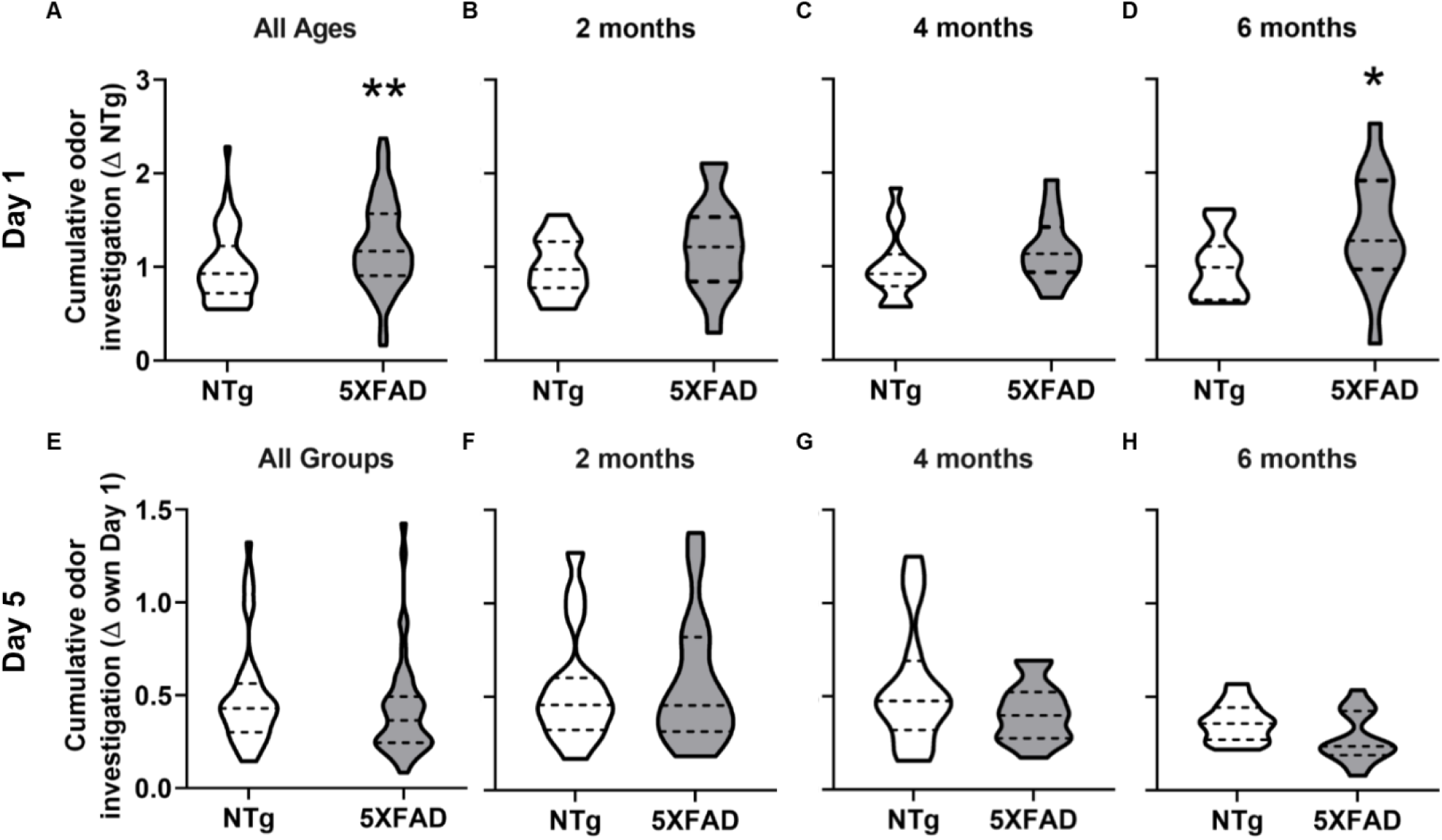
5XFAD mice investigate odors longer than controls on the first day of testing. Cumulative odor investigation duration in **A**, all mice (n=44-48/group), **B**, 2-months, **C**, 4-months, and **D**, 6-months old mice (n=15-16/group). Cumulative odor investigation on Day 5 of testing in **E**, all mice (n=45-46/group), **F**, 2-months, **G**, 4-months, and **H**, 6-months old mice (n=14-16/group). *p<0.05, **p<0.01 NTg vs. 5XFAD.

**Figure 2.**
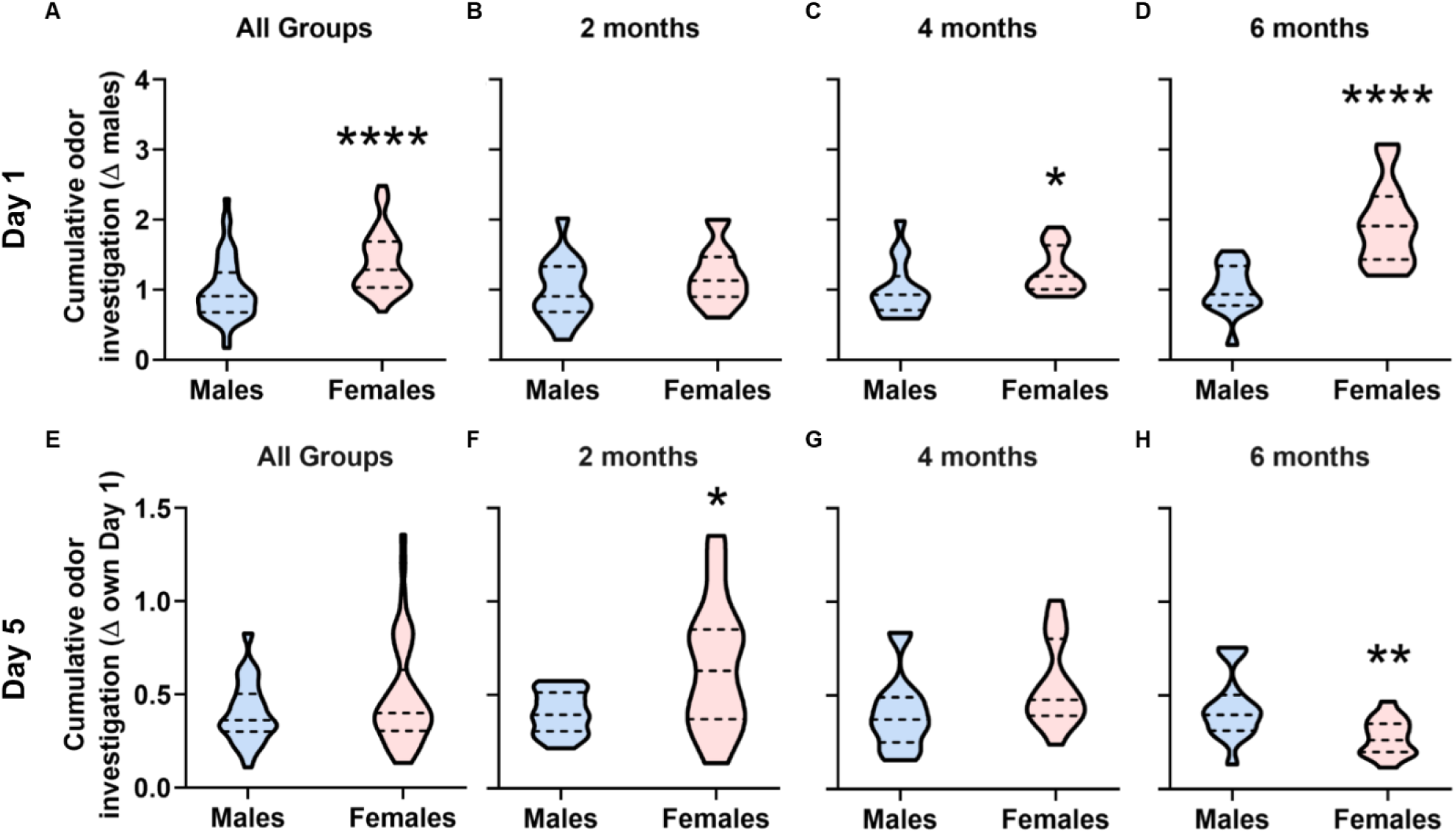
Females investigate odors longer than males on the first day of testing. Cumulative odor investigation duration in **A**, males and females (n=46-48/group), **B**, 2-months, **C**, 4-months, and **D**, 6-months old mice with genotypes collapsed (n=14-16/group). Cumulative odor investigation duration on Day 5 of testing in **E**, males and females (n=45-46/group), **F**, 2-months, **G**, 4-months, and **H**, 6-months old mice with genotypes collapsed (n=15-16/group). *p<0.05, **p<0.005, ****p<0.0001 vs. males.

## Results

### 5XFAD mice investigate odors longer than controls on the first testing day

We first analyzed the duration of odor investigation (cumulative, throughout the entire session) with data from all odors grouped, prior to determining if there was unique preference towards individual odors in later sections of the results. Examination of odor investigation durations revealed an effect of genotype (t(42)=2.98, p=0.005, collapsing all ages and sexes; **Figure 1A**) wherein 5XFAD mice exhibited longer odor investigation durations vs. NTg mice. We then separated the data in order to investigate the contributions of age upon the effect for 5XFAD mice to investigate odors longer. 6-months old 5XFAD mice engaged in odor investigation longer than age-matched NTg (t(13)=2.279, p=0.040; **Figures 1B – 1D**). There was no effect of genotype in 2-months and 4-months old mice (p>0.05). These results indicate that 5XFAD mutations and aging interact to influence odor investigation behavior.

While this study focuses on behavior during the first day of testing, we also examined the duration of odor investigation on testing day 5 to assess whether odor investigation habituated across testing days. Examination of odor investigation durations on day 5 revealed no effect of genotype with across all ages, nor within any specific age (p>0.05; **Figure 1E – 1H**). These results suggest 5XFAD mice have preserved odor habituation behavior.

### Females investigate odors longer than males on the first testing day

Due to established sex differences in AD (Nebel et al., 2018) and in pathological accumulation in this mouse model (Bundy et al., 2019), we sought to determine if sex influenced the above outcomes. Females exhibited longer odor investigation durations vs. males (t(92)=4.302, p<0.0001, collapsing ages and genotypes; **Figure 2A**). As above, we next separated the data to investigate the contributions of age upon the sex difference in odor investigation time. We found that 4- and 6-months old female mice exhibited longer odor investigation durations vs. age-matched males (t(29)=2.099, p=0.045; t(28)=5.314, p<0.0001, respectively, **Figure 2C & 2D**), with no sex difference at 2-months of age (p>0.05). These data show that females exhibit longer odor investigation durations vs. males, an effect carried by the older age groups.

As above, we investigated potential differences in odor habituation via assay of odor investigation duration on day 5 of testing. There was no effect of sex to influence odor habituation when collapsing across all ages and genotypes, (p<0.05; **Figure 2E**), although, within ages, 2-months old females exhibited longer odor investigation durations (t(29)=2.707, p=0.011; **Figure 2F**), while 6-months old females exhibited shorter odor investigation durations vs. age-matched males (t(29)=3.005, p=0.0048, **Figure 2H**), with no sex difference at 4-months of age (p>0.05). These data show that across-days odor habituation differs in females from 2- to 6-months of age. In all of the following sections we restrict our analyses to including only data from the first day of testing to reduce possible influences of across-days habituation.

### 5XFAD genotype impacts preferences towards individual odors

In all of the above analyses, odor investigation durations were collapsed across all odors as an assay for the tendencies of mice to investigate odors. However, collapsing data across all stimuli from a panel of molecularly diverse odors does not indicate changes towards a given odor which is essential in order to identify changes in odor preferences. Therefore, we next looked for possible influences of 5XFAD genotype on preferences towards specific odors.

As expected based upon the results in **Figure 1A**, this analysis revealed a main effect of genotype to influence duration of odor investigation across all age groups (F_(1, 456)_=20.07, p<0.0001, **Figure 3A**). 5XFAD mice exhibited a stronger preference for carvone and decanol vs. NTg control (t(456)=3.419, p=0.003; t(456)=3.233, p=0.007, respectively) with no effect for the other odors (p>0.05 for each).

**Figure 3.**
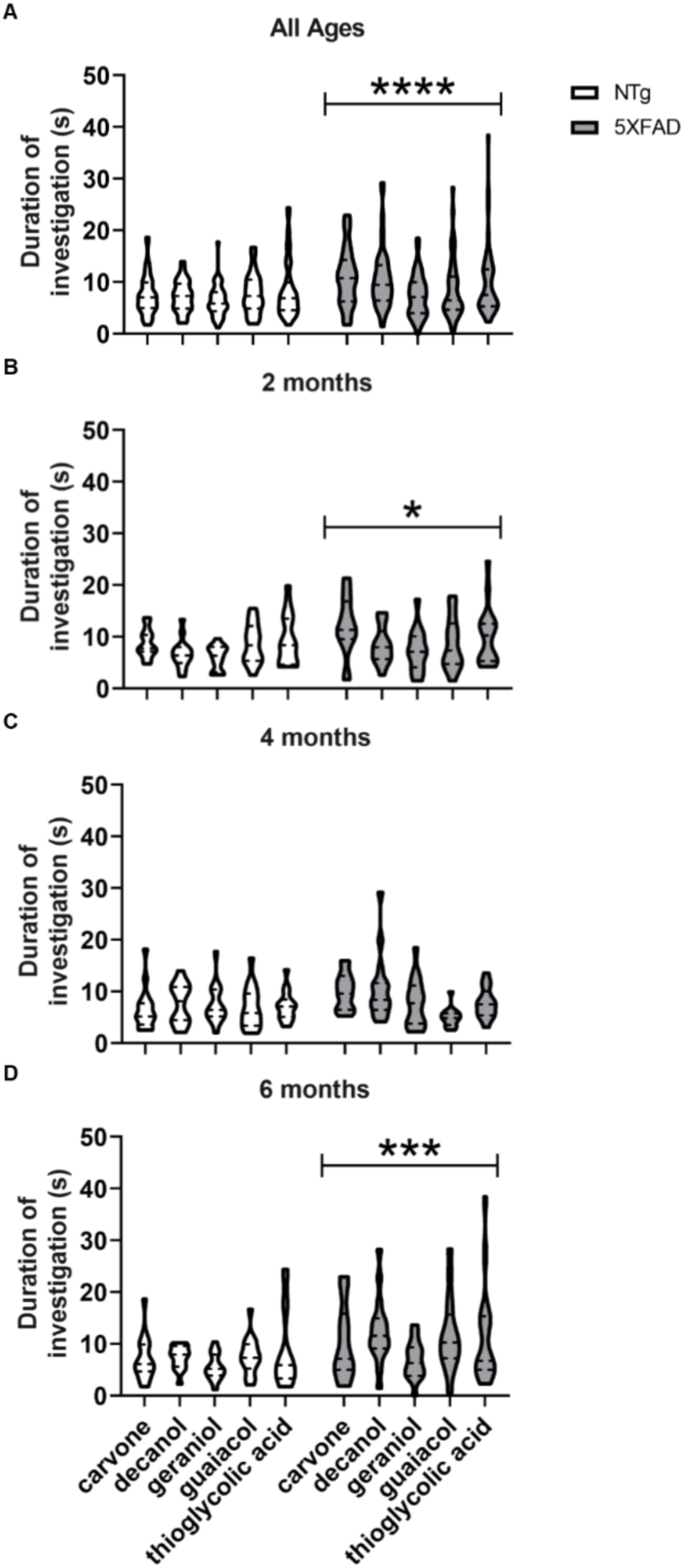
5XFAD genotype impacts preferences displayed towards individual odors. Duration of odor investigation in **A**, all mice (n=45-48/group), and in **B**, 2-months, **C**, 4-months, and **D**, 6-months old mice with sexes collapsed (n=15-16/group). *p<0.05, ***p<0.001, ****p<0.0001 vs. NTg.

When separating the data by age, we observed a main effect of genotype to influence duration of odor investigation in 2-months and 6-months mice (F_(1,146)_=4.92, p=0.028; F_(1,147)_=12.54, p<0.001, respectively) but not in 4-months mice (p>0.05). However, post hoc tests failed to uncover ‘attraction’ to an individual odor when testing within age groups for effects of genotype (p>0.05 for each).

### Sex and genotype impact preferences towards individual odors

While we found an effect of 5XFAD genotype on preferences towards odors, it is possible an effect was influenced by the pronounced sex difference wherein females exhibit longer odor investigation (as in **Figure 2**). Therefore, we next investigated if sex of the 5XFAD mice influences odor preferences.

There was an effect of sex to influence preferences for all odors (carvone: F_(1, 89)_=21.49, p<0.0001; decanol: F_(1, 89)_=10.02, p=0.002; geraniol: F_(1, 89)_=14.89, p<0.001; guaiacol: F_(1, 90)_=14.91, p<0.001; thioglycolic acid: F_(1, 88)_=7.895, p=0.006; **Figure 4)**. NTg females exhibited greater preferences vs. NTg males for some odors (carvone: t(89)=2.454, p=0.032; guaiacol: t(90)=2.496, p=0.029; p>0.05 for decanol, geraniol, and thioglycolic acid). Further, 5XFAD females exhibited greater preferences for most odors vs. 5XFAD males (carvone: t(89)=4.094, p<0.001; decanol: t(89)=3.323, p=0.003; geraniol: t(89)=4.017, p=0.002; guaiacol: t(90)=2.961, p=0.008; p>0.05 for thioglycolic acid). These data are in-line with our above findings from total odor investigation duration (**Figure 2**), where females investigated odors longer than males. The above analyses within individual odors indicates that sex influences odor preferences, as is well established in other mouse lines/strains (*e.g.*, (Baum & Keverne, 2002)).

**Figure 4.**
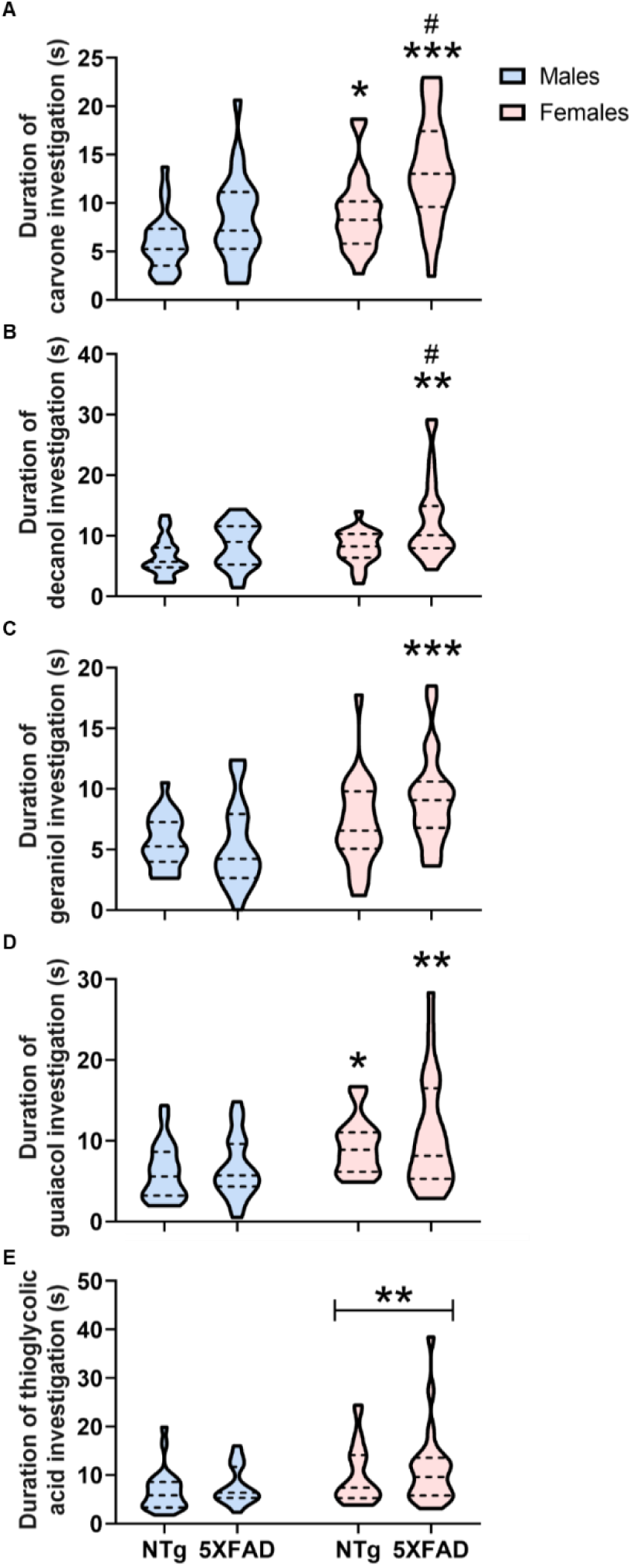
Sex and genotype impact preferences displayed towards individual odors. Odor investigation duration for **A**, carvone, **B**, decanol, **C**, geraniol, **D**, guaiacol, and **E**, thioglycolic acid (n=22-24/group). *p<.05, **p<.01, ***p<.001 vs. genotype-matched males. #p<.05 vs. NTg females.

5XFAD genotype also influenced preferences to some of the odors tested (carvone: F_(1, 89)_=14.79, p<0.001; decanol: F_(1, 89)_=12.3, p<0.001). Post hoc analysis revealed that the effect of genotype was carried by 5XFAD females, which displayed greater preferences vs. NTg females for carvone (t(89)=3.539, p=0.001) and decanol (t(89)=3.526, p=0.001; p>0.05 for geraniol, guaiacol, and thioglycolic acid). Overall, these data indicate that females within each age group exhibit more preference towards most odors vs. males. Further, in some cases, the enhanced preferences displayed by females is influenced by 5XFAD genotype.

As a final analysis into preference towards individual odors, we sought to test whether there was also an influence of age. Having uncovered that preference towards two of the odors in the panel was impacted by genotype and sex (**Figure 4A & 4B**), this resulted in the need of comparing possible effects of age not just within odor types, but also within genotypes (**Figure 5**).

**Figure 5.**
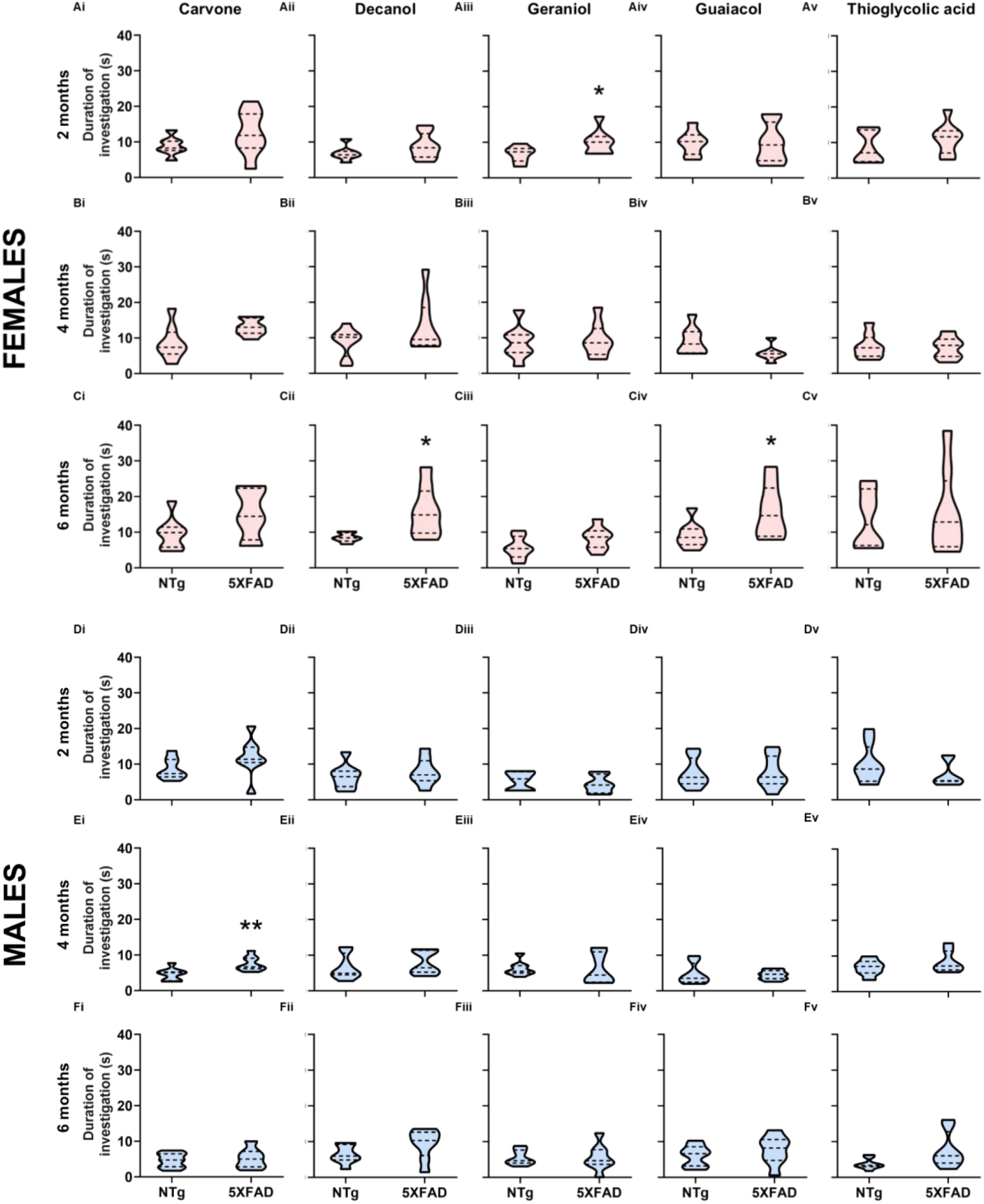
Sex, age, and genotype impact preferences displayed towards individual odors. Odor investigation duration for females at 2-months (**Ai-Av**), 4-months (**Bi-Bv**), and 6-months (**Ci-Cv**) and males at 2-months (**Di-Dv**), 4-months (**Ei-Ev**) and 6-months (**Fi-Fv**) for carvone, decanol, geraniol, guaiacol, and thioglycolic (n=7-8/group). *p<.05, **p<.01 vs. same-sex NTg counterpart.

This analysis revealed that within the 2-months females, there was an effect of genotype on preference for geraniol (t(13)=2.177, p=0.049) with no effect on any other odor (p>0.05 for all). In 6-months females, 5XFAD genotype influenced preference for decanol and guaiacol (t(13)=2.745, p=0.017; t(14)=2.29, p=0.038, respectively). There was no effect of genotype in 4-months females (p>0.05). Within males there was an effect of genotype wherein 5XFAD 4-months mice more greatly preferred carvone vs NTg (t(14)=3.135, p=0.007; p>0.05 for all other odors). There was no effect of genotype in 2-months and 6-months age groups (p>0.05 for all). These data indicate that females largely drive effects of 5XFAD mutations on odor investigation behaviors, particularly in the 6-months age group. Taken together, 5XFAD mice, especially females, show aberrant odor preferences.

### No effect of 5XFAD genotype on measures of general task engagement

Finally, we performed two analyses on the above data as simple ‘controls’ to indicate the mice, regardless of group, were capable of engaging in the task. More greatly aged 5XFAD mice are known to have impaired motor function (O’Leary et al., 2018, 2020) and changes in the ability of mice to nose-poke would confound use of this measure as a variable whereby to assay odor investigation and preferences. We first examined inter-poke interval, which is indicative of an animal’s ability to coordinate motor movements from one odor port to another. There was neither an effect of genotype nor sex to influence inter-poke intervals in both 2- and 4-month old mice (p>0.05 each; **Figs 6A & 6B**). In 6-month old mice there is an effect of sex (F_(1, 26)_=5.307, p=0.03; **Figure 6C**), but not genotype (p>0.05), to shorten inter-poke interval.

**Figure 6.**
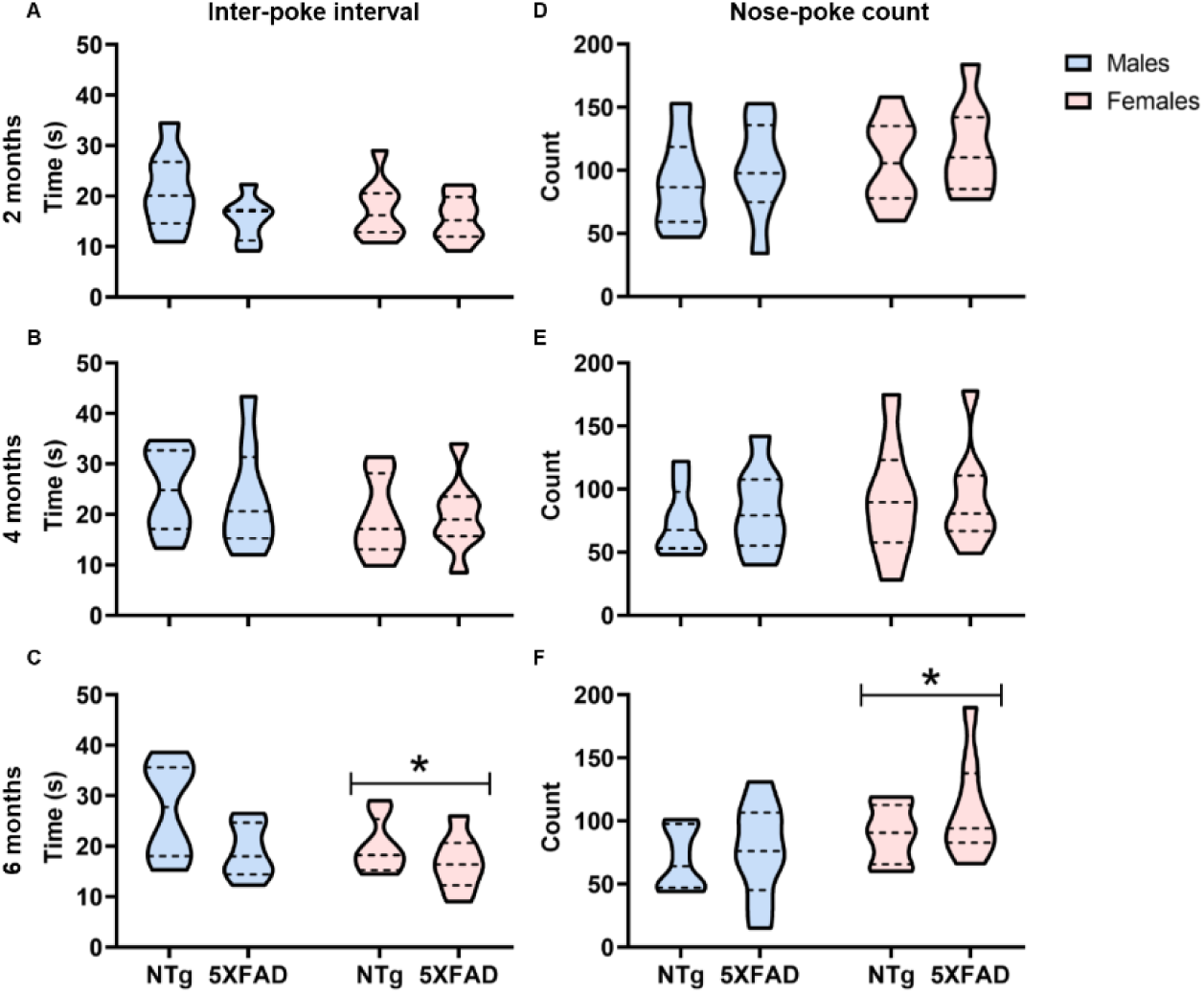
No effect of 5XFAD genotype on measures of general task engagement. Inter-poke interval in **A**, 2-months (n=7-8/group), **B**, 4-months (n=7-8/group), and **C**, 6-months old mice (n=8/group). Nose-poke count in **D**, 2-months (n=8/group), **E**, 4-months (n=8/group), and **F**, 6-months old mice (n=8/group). *p<.05 vs. males.

An additional index of general task engagement is the total number of nose-pokes animals execute within a session. There was neither an effect of genotype nor sex to influence nose-poke count in both 2- and 4-months old mice (p>0.05 each; **Figure 6D & 6E**). 6-months old mice exhibited a significant effect of sex (F_(1, 28)_=5.212, p=0.03; **Figure 6F**) but not genotype (p>0.05), to increase nose-poke count. Reconciling these outcomes, these significant differences however do not support impairments in the task. Indeed, a greater number of pokes in 6-months old female 5XFAD mice (**Figure 6C**), together with their reduced inter-poke intervals (**Figure 6F**), point towards increased task engagement. Taken together, data from both analyses suggest the differences in odor investigation and preferences observed in the present study are not due to deficits in task engagement.

## Discussion

We used the 5XFAD mouse AD model in order to define odor investigation behavior using the five-port odor multiple-choice task. Our long-term goal is to understand how odor hedonics and possibly other affective symptoms of AD result mechanistically. To this end, we assessed if odor preferences, reflected by the time sampling individual odors, are perturbed in the 5XFAD mouse AD model. We found, paradoxical to what is known in persons with AD to display reduced hedonic ratings of odors (Joussain et al., 2015), that 5XFAD mice sample certain odors longer, reflective of increased hedonics.

### No gross behavioral impairments are detected in 5XFAD mice in the 5-port odor multiple choice task

As an important starting point, our results indicate that 5XFAD mice do not display overt impairments in their ability to engage in the task we used (**Figure 6**). This suggests that 5XFAD mice, at the ages we tested, have intact motor function at least as it is required to coordinate nose-pokes in this task. A robust literature suggests that there are significant motor impairments in these mice occurring at much later ages (O’Leary et al., 2018, 2020). If anything, female mice in our most aged group (6-months) showed changes in task engagement measures suggesting *increased* engagement in the task (engaged more frequently and promptly between pokes; **Figure 6**), yet this was not influenced by genotype. In addition to task engagement, future studies should include a comprehensive assay of mood-associated and locomotor behaviors as this may influence performance and/or engagement in behavioral tests. This is especially important given that motor impairments and alterations in indices of anxiety-like behaviors have been reported in 5XFAD mice (Braun & Feinstein, 2019; Jawhar et al., 2010), the latter of which may be supported by altered social behaviors and sensory processing (Flanigan et al., 2014).

We also uncovered that 2-6-months old 5XFAD mice of both sexes are capable of engaging with odors (**Figure 1**). This latter outcome suggests the 5XFAD mice do not have grossly impaired abilities to detect and discriminate among odors since if they would simply generalize across all odors, one would predict to display similar investigation regardless of the odor. In contrast, our analyses among the individual odors (**Figures 4 & 5**) support that mice can differentiate among the odors we used. These results at their most basic level are consistent with a report from one group that measured odor detection thresholds using sophisticated olfactometry in 5XFAD mice (Roddick et al., 2016). Using the go/no-go olfactometer as an assay, Roddick and colleagues quantified odor detection thresholds and concluded that odor sensitivity is not dramatically impaired in 6-months old 5XFAD mice (Roddick et al., 2016). A study by a separate group used the buried food pellet test to perform testing of odor-influenced food motivation behavior (Xiao et al., 2015). The work by Xiao indicated possible deficits in food-finding behavior at three months of age which may suggest deficits in olfaction. However, this test is difficult to use as a specific metric of olfaction since it relies upon the ability for animals to coordinate foraging behavior which is fundamentally regulated by states of motivation. Overall, the results of the present study together with especially that of Roddick et al., (2016) highlight that the most basic aspects of olfaction are largely preserved in 5XFAD mice. This outcome is perhaps surprising given that the brains of 6-months old 5XFAD mice are burdened with extensive AD pathologies (Devi & Ohno, 2016; Oakley et al., 2006), including within the olfactory bulbs themselves (Oakley et al., 2006; Wang et al., 2017; Xiao et al., 2015).

### Alterations in odor investigation behavior and odor preferences in 5XFAD mice

Collapsing across all odors, our results demonstrate that 5XFAD mice, all ages and both sexes included, investigate odors longer than NTg controls (**Figure 1**). This was due to longer investigation behavior specifically in the 6-month old animals, but not the 2- and 4-months mice. We used these data pooled across all odors as a simple metric of generalized odor investigation behavior. Importantly, generalized odor investigation behavior may not be used to understand hedonics necessarily, since this may also be resultant from enhanced arousal and/or novelty behavior. Nevertheless, it is informative to consider the tendency for 5XFAD mice, especially 6-months old female 5XFAD mice, to display enhanced propensities to investigate odors.

By analyzing the data within individual odors grouped by sex and age, we uncovered several examples supporting our conclusion that 5XFAD mice display altered hedonics to certain odors. Two of the five odors employed (carvone and decanol) often entailed aberrant investigation responses in 5XFAD mice, but did so depending on age and sex. Notably, we anticipate the uncovered changes in odor hedonics of the 5XFAD mice may be more prevalent than that found herein. Our use of a simple panel of five odors was selected to provide a spectrum of odor/chemical types to thereby yield a first-pass screen of olfactory preferences in 5XFAD mice. Due the extensive numbers of animals tested in this study, and testing labor this study required, we did not also include additional panels of odors to further assay olfactory abilities. Certainly, investigating the behavior of 5XFAD mice in the 5-port task with differing panels of odors is an important future goal to provide a complete assessment of the magnitude of odor preference alterations. One reason for this is that the differences in preferences we found were not highly overt like one would expect to see if a mouse was allowed to investigate conspecific urine or a food source odor versus that of a predator. Instead, the preferences were subtle and suggest modest, although functionally significant at a behavioral level, changes in affective brain systems supporting odor preferences. In the future it will be important to understand what brain systems might be impacted in these animals which results in altered preferences to select odors.

Perplexingly, while in persons with AD, hedonic ratings towards odors are notably reduced (Joussain et al., 2015), we found that 5XFAD mice show opposite responses when engaging with odors during altered hedonic states. Indeed, our results show that in some cases, 5XFAD mice investigate odors longer, and specific odors longer, than control mice (**Figures 4 & 5**). While increases in odor investigation are known to be reliable measures of odor preferences in both humans and mice (Mandairon et al., 2009), why 5XFAD mice would display increased hedonic responses to odors is unclear. It is possible that an AD pathologically-burdened mouse model displays hedonic responses to odors in fundamentally different manners than a human.

### Sex influences on odor investigation and preferences

Olfaction is influenced by sex and gonadal hormones (Dillon et al., 2013; Doty & Cameron, 2009; Good et al., 1976; Kass et al., 2017; Sorwell et al., 2008; Wesson et al., 2006). This also holds true to odor preferences (Baum & Keverne, 2002; Beach, 1974). In line with this, we found that females exhibited longer durations of odor investigation compared to males specifically at 4- and 6-months of age (**Figure 2**). We also found that females across all ages and in both genotypes exhibited higher preference for most of the odors as compared to males (**Figure 4**). Further, while the 5-port apparatus was not designed to assay odor habituation since mice are required to nose poke to obtain the odor, the 6-months old female mice habituated to odors more greatly (at day 5) compared to males (**Figure 2**).

AD differentially impacts males and females (Nebel et al., 2018), and this is somewhat recapitulated in neuronal pathology in the 5XFAD mouse model wherein females exhibit more AD-associated pathology compared to males (Bundy et al., 2019). Indeed, the longer odor investigation duration of the 5XFAD mice upon day 1 was driven primarily by the behavior of the females. Consistent with this, male and female 5XFAD mice display different odor learning curves, yet comparable odor detection abilities (Roddick et al., 2016). Taken together, these results emphasize that it is important to consider sex as a biological variable in AD model research, especially given the established sex differences in human AD presentation, development, and progression (Nebel et al., 2018).

### Conclusions

Our results show that young adult 5XFAD mice investigate odors longer than NTg controls. This effect was not simply due to genotype. Indeed, prolonged odor investigation in 5XFAD mice was influenced significantly by both age and sex with females investigating odors longer than males regardless of genotype. These results provide some evidence for altered odor investigation behaviors and further, odor preferences, in an AD mouse model, the latter of which was surprisingly increased among 5XFAD mice towards some odors. Future work employing larger odor sets are needed to best define the nature of these changes in preferences, as are studies to explore their mechanistic causes.

## Acknowledgements

This work was supported by National Institutes of Health grants: NIDCD R01DC016519 (to D.W. and P.B.) and NIDCD R01DC014443 to D.W. (the latter including a supplement from the NIA to support research into Alzheimer’s disease to P.B. and D.W.) and NIDA R01DA049545 and R01DA049449 (to D.W.).

## References

Baum, M. J., & Keverne, E. B. (2002). Sex difference in attraction thresholds for volatile odors from male and estrous female mouse urine. Hormones and Behavior, 41(2), 213–219.

Beach, F. A. (1974). Effects of gonadal hormones on urinary behavior in dogs. Physiology and Behavior, 12(6), 1005–1013.

Braun, D., & Feinstein, D. L. (2019). The locus coeruleus neuroprotective drug vindeburnol normalizes behavior in the 5xFAD transgenic mouse model of Alzheimer’s disease. Brain Research, 1702, 29–37. https://doi.org/https://doi.org/10.1016/j.brainres.2017.12.028

Bundy, J. L., Vied, C., Badger, C., & Nowakowski, R. S. (2019). Sex-biased hippocampal pathology in the 5XFAD mouse model of Alzheimer’s disease: A multi-omic analysis. The Journal of Comparative Neurology, 527(2), 462–475. https://doi.org/10.1002/cne.24551

Cheng, N., Cai, H., & Belluscio, L. (2011). In Vivo Olfactory Model of APP-Induced Neurodegeneration Reveals a Reversible Cell-Autonomous Function. The Journal of Neuroscience, 31(39), 13699–13704. https://doi.org/10.1523/jneurosci.1714-11.2011

Devanand, D. P., Liu, X., Tabert, M. H., Pradhaban, G., Cuasay, K., Bell, K., de Leon, M. J., Doty, R. L., Stern, Y., & Pelton, G. H. (2008). Combining Early Markers Strongly Predicts Conversion from Mild Cognitive Impairment to Alzheimer’s Disease. Biological Psychiatry, 64(10), 871–879. http://www.sciencedirect.com/science/article/B6T4S-4T8TW52-1/2/166fa8cd739a0242ecc9096da4f9e303

Devi, L., & Ohno, M. (2016). Cognitive benefits of memantine in Alzheimer’s 5XFAD model mice decline during advanced disease stages. Pharmacology Biochemistry and Behavior, 144, 60–66. https://doi.org/https://doi.org/10.1016/j.pbb.2016.03.002

Dillon, T. S., Fox, L. C., Han, C., & Linster, C. (2013). 17β-estradiol enhances memory duration in the main olfactory bulb in CD-1 mice. Behavioral Neuroscience, 127(6), 923–931. https://doi.org/10.1037/a0034839

Doty, R. L. (2017). Olfactory dysfunction in neurodegenerative diseases: is there a common pathological substrate? The Lancet Neurology, 16(6), 478–488. https://doi.org/https://doi.org/10.1016/S1474-4422(17)30123-0

Doty, R. L., & Cameron, E. L. (2009). Sex differences and reproductive hormone influences on human odor perception. Physiology & Behavior, 97(2), 213–228. https://doi.org/10.1016/j.physbeh.2009.02.032

Flanigan, T. J., Xue, Y., Kishan Rao, S., Dhanushkodi, A., & McDonald, M. P. (2014). Abnormal vibrissa-related behavior and loss of barrel field inhibitory neurons in 5xFAD transgenics. Genes, Brain, and Behavior, 13(5), 488–500. https://doi.org/10.1111/gbb.12133

Good, P. R., Geary, N., & Engen, T. (1976). The effect of estrogen on odor detection. Chemical Senses and Flavor, 2, 45–50.

Gopinath, B., Anstey, K. J., Kifley, A., & Mitchell, P. (2012). Olfactory impairment is associated with functional disability and reduced independence among older adults. Maturitas, 72(1), 50–55. https://doi.org/10.1016/j.maturitas.2012.01.009

Jagetia, S., Milton, A. J., Stetzik, L. A., Liu, S., Pai, K., Arakawa, K., Mandairon, N., & Wesson, D. W. (2018). Inter- and intra-mouse variability in odor preferences revealed in an olfactory multiple-choice test. In Behavioral Neuroscience (Vol. 132, Issue 2, pp. 88–98). American Psychological Association. https://doi.org/10.1037/bne0000233

Jawhar, S., Trawicka, A., Jenneckens, C., Bayer, T. A., & Wirths, O. (2010). Motor deficits, neuron loss, and reduced anxiety coinciding with axonal degeneration and intraneuronal A[beta] aggregation in the 5XFAD mouse model of Alzheimer’s disease. Neurobiology of Aging.

Joussain, P., Bessy, M., Fournel, A., Ferdenzi, C., Rouby, C., Delphin-Combe, F., Krolak-Salmon, P., & Bensafi, M. (2015). Altered Affective Evaluations of Smells in Alzheimer’s Disease. Journal of Alzheimer’s Disease, 49(2), 433–441.

Kass, M. D., Czarnecki, L. A., Moberly, A. H., & McGann, J. P. (2017). Differences in peripheral sensory input to the olfactory bulb between male and female mice. Scientific Reports, 7(1), 45851. https://doi.org/10.1038/srep45851

Kimura, R., & Ohno, M. (2009). Impairments in remote memory stabilization precede hippocampal synaptic and cognitive failures in 5XFAD Alzheimer mouse model. Neurobiology of Disease, 33(2), 229–235. https://doi.org/https://doi.org/10.1016/j.nbd.2008.10.006

Mandairon, N., Poncelet, J., Bensafi, M., & Didier, A. (2009). Humans and Mice Express Similar Olfactory Preferences. PLoS ONE, 4(1), e4209. https://doi.org/10.1371/journal.pone.0004209

Morales-Corraliza, J., Schmidt, S. D., Mazzella, M. J., Berger, J. D., Wilson, D. A., Wesson, D. W., Jucker, M., Levy, E., Nixon, R. A., & Mathews, P. M. (2012). Immunization targeting a minor plaque constituent clears β-amyloid and rescues behavioral deficits in an Alzheimer’s disease mouse model. Neurobiology of Aging, in press(0). https://doi.org/10.1016/j.neurobiolaging.2012.04.007

Murphy, C. (2008). The chemical senses and nutrition in older adults. Journal of Nutrition For the Elderly, 27(3–4), 247–265.

Murphy, C. (2019). Olfactory and other sensory impairments in Alzheimer disease. Nature Reviews Neurology, 15(1), 11–24. https://doi.org/10.1038/s41582-018-0097-5

Nebel, R. A., Aggarwal, N. T., Barnes, L. L., Gallagher, A., Goldstein, J. M., Kantarci, K., Mallampalli, M. P., Mormino, E. C., Scott, L., Yu, W. H., Maki, P. M., & Mielke, M. M. (2018). Understanding the impact of sex and gender in Alzheimer’s disease: A call to action. Alzheimer’s & Dementia : The Journal of the Alzheimer’s Association, 14(9), 1171–1183. https://doi.org/10.1016/j.jalz.2018.04.008

O’Leary, T., Mantolino, H., Stover, K., & Brown, R. (2020). Age-related deterioration of motor function in male and female 5xFAD mice from 3 to 16 months of age. Genes Brain and Behavior, 19(3), e12538. https://doi.org/10.1111/gbb.12538

O’Leary, T., Robertson, A., Chipman, P. H., Rafuse, V. F., & Brown, R. E. (2018). Motor function deficits in the 12 month-old female 5xFAD mouse model of Alzheimer’s disease. Behavioural Brain Research, 337, 256–263. https://doi.org/10.1016/j.bbr.2017.09.009

Oakley, H., Cole, S. L., Logan, S., Maus, E., Shao, P., Craft, J., Guillozet-Bongaarts, A., Ohno, M., Disterhoft, J., Van Eldik, L., Berry, R., & Vassar, R. (2006). Intraneuronal beta-Amyloid Aggregates, Neurodegeneration, and Neuron Loss in Transgenic Mice with Five Familial Alzheimer’s Disease Mutations: Potential Factors in Amyloid Plaque Formation. Journal of Neuroscience, 26(40), 10129–10140. https://doi.org/10.1523/jneurosci.1202-06.2006

Rey, N. L., Wesson, D. W., & Brundin, P. (2018). The olfactory bulb as the entry site for prion-like propagation in neurodegenerative diseases. Neurobiology of Disease, 109, 226–248. https://doi.org/http://dx.doi.org/10.1016/j.nbd.2016.12.013

Roddick, K. M., Roberts, A. D., Schellinck, H. M., & Brown, R. E. (2016). Sex and Genotype Differences in Odor Detection in the 3×Tg-AD and 5XFAD Mouse Models of Alzheimer’s Disease at 6 Months of Age. Chemical Senses, 41(5), 433–440. https://doi.org/10.1093/chemse/bjw018

Sorwell, K. G., Wesson, D. W., & Baum, M. J. (2008). Sexually dimorphic enhancement by estradiol of male urinary odor detection thresholds in mice. Behavioral Neuroscience, 122(4), 788–793. https://doi.org/2008-09788-007 [pii] 10.1037/0735-7044.122.4.788

Wang, Y., Wu, Z., Bai, Y.-T., Wu, G.-Y., & Chen, G. (2017). Gad67 haploinsufficiency reduces amyloid pathology and rescues olfactory memory deficits in a mouse model of Alzheimer’s disease. Molecular Neurodegeneration, 12(1), 73. https://doi.org/10.1186/s13024-017-0213-9

Wesson, D. W., Borkowski, A. H., Landreth, G. E., Nixon, R. A., Levy, E., & Wilson, D. A. (2011). Sensory Network Dysfunction, Behavioral Impairments, and Their Reversibility in an Alzheimer’s beta-Amyloidosis Mouse Model. The Journal of Neuroscience, 31(44), 15962–15971. https://doi.org/10.1523/jneurosci.2085-11.2011

Wesson, D. W., Keller, M., Douhard, Q., Baum, M. J., & Bakker, J. (2006). Enhanced urinary odor discrimination in female aromatase knockout (ArKO) mice. Hormones and Behavior, 49(5), 580–586.

Wesson, D. W., Levy, E., Nixon, R. A., & Wilson, D. A. (2010). Olfactory Dysfunction Correlates with β-Amyloid Plaque Burden in an Alzheimer’s Disease Mouse Model. The Journal of Neuroscience, 30(2), 505–514.

Xiao, N.-A., Zhang, J., Zhou, M., Wei, Z., Wu, X.-L., Dai, X.-M., Zhu, Y.-G., & Chen, X.-C. (2015). Reduction of Glucose Metabolism in Olfactory Bulb is an Earlier Alzheimer’s Disease-related Biomarker in 5XFAD Mice. Chinese Medical Journal, 128(16), 2220–2227. https://doi.org/10.4103/0366-6999.162507

Xu, W., Fitzgerald, S., Nixon, R. A., Levy, E., & Wilson, D. A. (2015). Early hyperactivity in lateral entorhinal cortex is associated with elevated levels of AβPP metabolites in the Tg2576 mouse model of Alzheimer’s disease. Experimental Neurology, 264, 82–91. https://doi.org/https://doi.org/10.1016/j.expneurol.2014.12.008

Yoo, S.-J., Lee, J.-H., Kim, S. Y., Son, G., Kim, J. Y., Cho, B., Yu, S.-W., Chang, K.-A., Suh, Y.-H., & Moon, C. (2017). Differential spatial expression of peripheral olfactory neuron-derived BACE1 induces olfactory impairment by region-specific accumulation of β-amyloid oligomer. Cell Death Disease, 8, e2977. https://doi.org/10.1038/cddis.2017.349

